# Temporal changes in gut microbiota and signaling molecules of the gut–brain axis in mice fed meat protein diets

**DOI:** 10.1101/329953

**Authors:** Yunting Xie, Guanghong Zhou, Chao Wang, Xinglian Xu, Chunbao Li

## Abstract

The purpose of this study was to characterize the dynamical changes of gut microbiota and explore the influence on bidirectional communications between the gut and the brain during a relatively long-term intake of different protein diets. The C57BL/6J mice were fed casein, soy protein and four kinds of processed meat proteins at a normal dose of 20% for 8 months. Protein diets dramatically affected the microbial composition and function and also the signaling molecule levels of the gut–brain axis in a dynamic manner, which consequently affected growth performance. *Alistipes*, Clostridiales vadinBB60, *Anaerotruncus*, *Blautia* and *Oscillibacter* had a relatively fast response to the diet, while Bacteroidales S24-7, *Ruminiclostridium*, *Ruminococcaceae UCG-014*, *Coriobacteriaceae UCG-002* and *Bilophila* responded slowly. *Rikenellaceae RC9 gut*, *Faecalibaculum* and Lachnospiraceae showed a continuous change with feeding time. Bacteroidales S24-7 abundance increased from 4 months to 8 months, whereas those of *Rikenellaceae RC9 gut, Akkermansia, Alistipes* and *Anaerotruncus* remarkably decreased. Five and fifteen biological functions of microbiota were affected at 4 months and 8 months, respectively, and sixteen functions were observed to change over feeding time. Moreover, 28 and 48 specific operational taxonomy units were associated with the regulation of serotonin, peptide YY, leptin and insulin levels at two time points. Ruminococcaceae was positively associated with Lachnospiraceae and negatively associated with Bacteroidales S24-7. These results give an important insight into the effect of gut microbiota on the bidirectional communications between the gut and the brain under a certain type of diet.

**Importance:** Many gastrointestinal and neuropsychiatric disorders may have a common pathophysiologic mechanism, involving bidirectional brain–gut axis signaling through humoral and neural pathways. The gut microbiota plays an important role in the communications between the gut and the brain. Recent evidence suggests that a growing number of subjects suffer from the above disorders. The significance of this study lies in the finding that long-term intake of different proteins at a normal dose induces dynamic alterations of specific microbiota in mice, which consequently affect bidirectional communications between the gut and the brain and results in different growth performance through dynamically regulating signaling molecule levels. Furthermore, this study indicates that intake of the same diet for a long time, irrespective of the diet source, may have an adverse effect on host health by altering gut microbiota.

## Introduction

In recent years, the gut–brain axis has attracted great interest, and previous studies have shown that the gut microbiota plays an important role in the bidirectional communications between the gut and the brain, coined the microbiota–gut–brain axis (1). The brain ensures proper maintenance and coordination of gastrointestinal functions. In turn, the gut microbiota has a great influence on central nervous system activities and host behavior, with chemical signaling of the gut–brain axis being involved. The trillions of microbes in the gastrointestinal tract are considered a complex and dynamic ecosystem that has coevolved with the host (2). Many factors have a certain impact on gut microbiota, e.g., genetics, geographic origin, age, medication and diet (3, 4), among which diet is the dominant modulator of the composition and function of gut microbiota (5). The majority of dietary proteins are digested into peptides and free amino acids in the small intestine, but some proteins cannot be digested and absorbed in the small intestine, so these enter the large intestine for microbial fermentation (6, 7). High-protein diets have been shown to alter the gut microbial composition (8, 9). The temporal microbial changes were also observed in feces after 6 weeks of a high-protein diet intake (10). Moreover, some studies indicated that dietary protein sources affect the gut microbial composition (11, 12).

Meat is known to be an important source of high-quality protein that contains all essential amino acids. In processed meat, the processing methods may lead to different degrees of protein oxidation and denaturation and hence cause protein aggregation and changes in secondary structures (13, 14). Moderate denaturation will increase the degradation of meat proteins, but various amino acid modifications might lead to the formation of “limit peptides,” which are not further broken down and thus result in a reduction of protein bioavailability (15, 16). Our *in vitro* studies showed that protein digestibility and digested products differed among cooked pork, emulsion-type sausage, dry-cured pork and stewed pork (17). Most studies have focused on the short-term effects of dietary proteins, and few data are available on the temporal variations in gut microbial composition. This study aimed to investigate whether a relatively long-term intake of proteins from processed meat affects the gut microbiota and the bidirectional communications between the gut and the brain.

## Results

### Composition and functions of gut microbiota

#### Richness and diversity

A total of 1801422 reads were obtained from all fecal samples with an average of 31059 reads per sample. Using the identification criterion of 97% sequence similarity at the species level, a total of 15741 operation taxonomy units (OTUs) were identified from all the samples, with an average of 271 OTUs per sample. The rarefaction and Shannon– Wiener curves tend to approach the saturation plateau and the Good’s coverage indices were greater than 99%, indicating sufficient data sampling and adequate sequencing depth.

Community richness estimators (Chao and ACE), and diversity indices (Shannon and Simpson) were calculated in order to evaluate the alpha diversity (Table 1). The protein diets significantly affected ACE and Chao values at both time points. The ACE and Chao values of the casein diet (CD) and stewed pork protein diet (SPPD) groups were significantly lower than those of other groups at 4 months, while the values were the highest for the soy protein diet (SPD) group at 8 months. The Shannon and Simpson values were not affected by diets at 4 months, but the Shannon value of the SPD group was higher than other groups at 8 months. In addition, the Shannon values decreased with feeding time, indicating that the microbial diversity may be reduced during long-term feeding of the same diet.

**Table 1.**
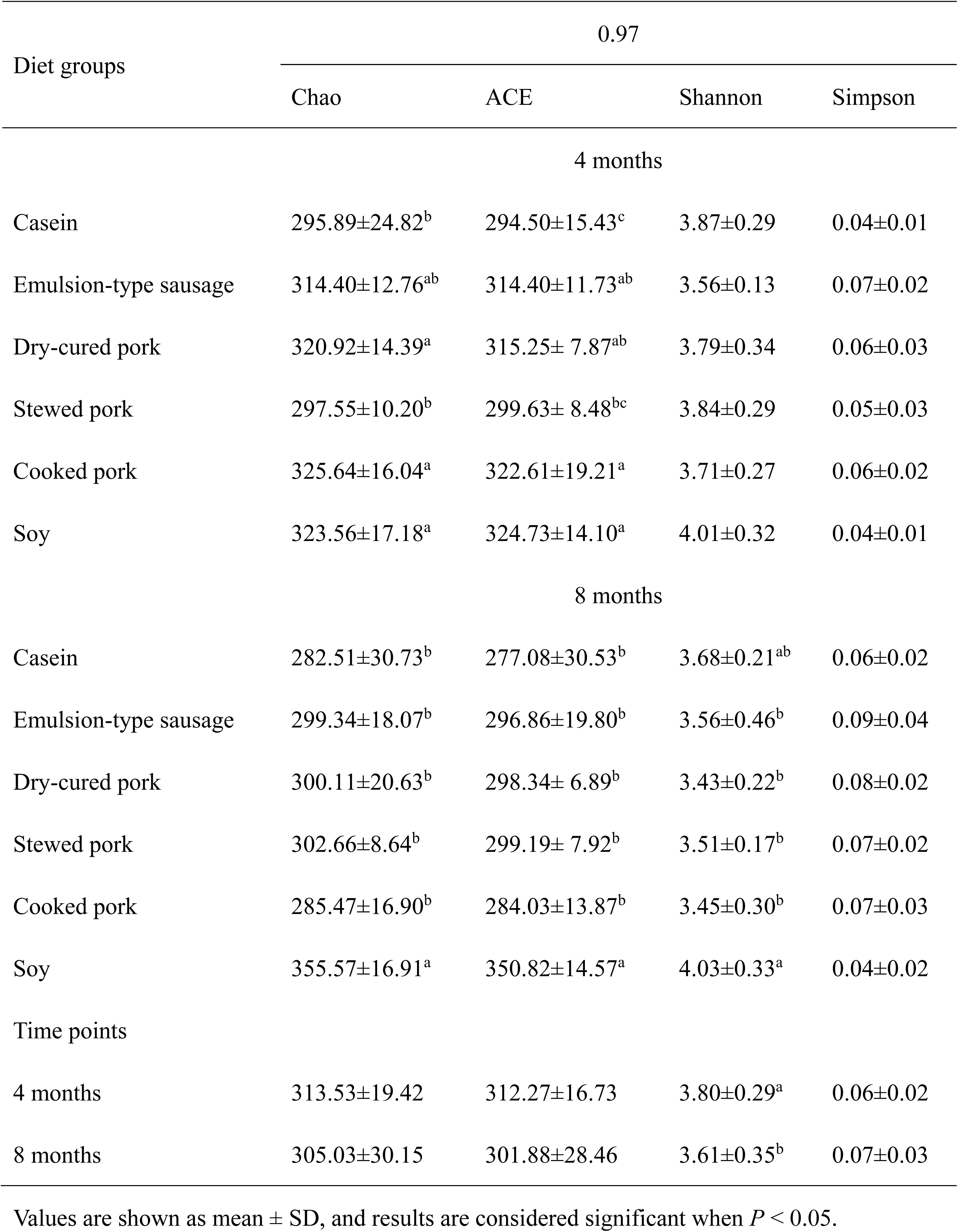
Richness and diversity indices of fecal microbiota. Values are shown as mean ± SD, and results are considered significant when *P* < 0.05.

| Diet groups | 0.97 |  |  |  |
| --- | --- | --- | --- | --- |
|  | Chao | ACE | Shannon | Simpson |
| 4 months |  |  |  |  |
| Casein | 295.89±24.82<sup>b</sup> | 294.50±15.43<sup>c</sup> | 3.87±0.29 | 0.04±0.01 |
| Emulsion-type sausage | 314.40±12.76<sup>ab</sup> | 314.40±11.73<sup>ab</sup> | 3.56±0.13 | 0.07±0.02 |
| Dry-cured pork | 320.92±14.39<sup>a</sup> | 315.25± 7.87<sup>ab</sup> | 3.79±0.34 | 0.06±0.03 |
| Stewed pork | 297.55±10.20<sup>b</sup> | 299.63± 8.48<sup>bc</sup> | 3.84±0.29 | 0.05±0.03 |
| Cooked pork | 325.64±16.04<sup>a</sup> | 322.61±19.21<sup>a</sup> | 3.71±0.27 | 0.06±0.02 |
| Soy | 323.56±17.18<sup>a</sup> | 324.73±14.10<sup>a</sup> | 4.01±0.32 | 0.04±0.01 |
| 8 months |  |  |  |  |
| Casein | 282.51±30.73<sup>b</sup> | 277.08±30.53<sup>b</sup> | 3.68±0.21<sup>ab</sup> | 0.06±0.02 |
| Emulsion-type sausage | 299.34±18.07<sup>b</sup> | 296.86±19.80<sup>b</sup> | 3.56±0.46<sup>b</sup> | 0.09±0.04 |
| Dry-cured pork | 300.11±20.63<sup>b</sup> | 298.34± 6.89<sup>b</sup> | 3.43±0.22<sup>b</sup> | 0.08±0.02 |
| Stewed pork | 302.66±8.64<sup>b</sup> | 299.19± 7.92<sup>b</sup> | 3.51±0.17<sup>b</sup> | 0.07±0.02 |
| Cooked pork | 285.47±16.90<sup>b</sup> | 284.03±13.87<sup>b</sup> | 3.45±0.30<sup>b</sup> | 0.07±0.03 |
| Soy | 355.57±16.91<sup>a</sup> | 350.82±14.57<sup>a</sup> | 4.03±0.33<sup>a</sup> | 0.04±0.02 |
| Time points |  |  |  |  |
| 4 months | 313.53±19.42 | 312.27±16.73 | 3.80±0.29<sup>a</sup> | 0.06±0.02 |
| 8 months | 305.03±30.15 | 301.88±28.46 | 3.61±0.35<sup>b</sup> | 0.07±0.03 |

Principal coordinate analysis (PCoA) on the OTU level confirmed that the fecal microbiota in the CD, SPD and emulsion-type sausage protein diet (ESPD) was distinct from that in other meat protein diet groups at 4 months. Diet groups were also well separated at 8 months, except the cooked pork protein diet (CPPD) and SPPD groups. In addition, the fecal microbiota was well separated between the two time points (Fig. 1A to C).

**Figure 1.**
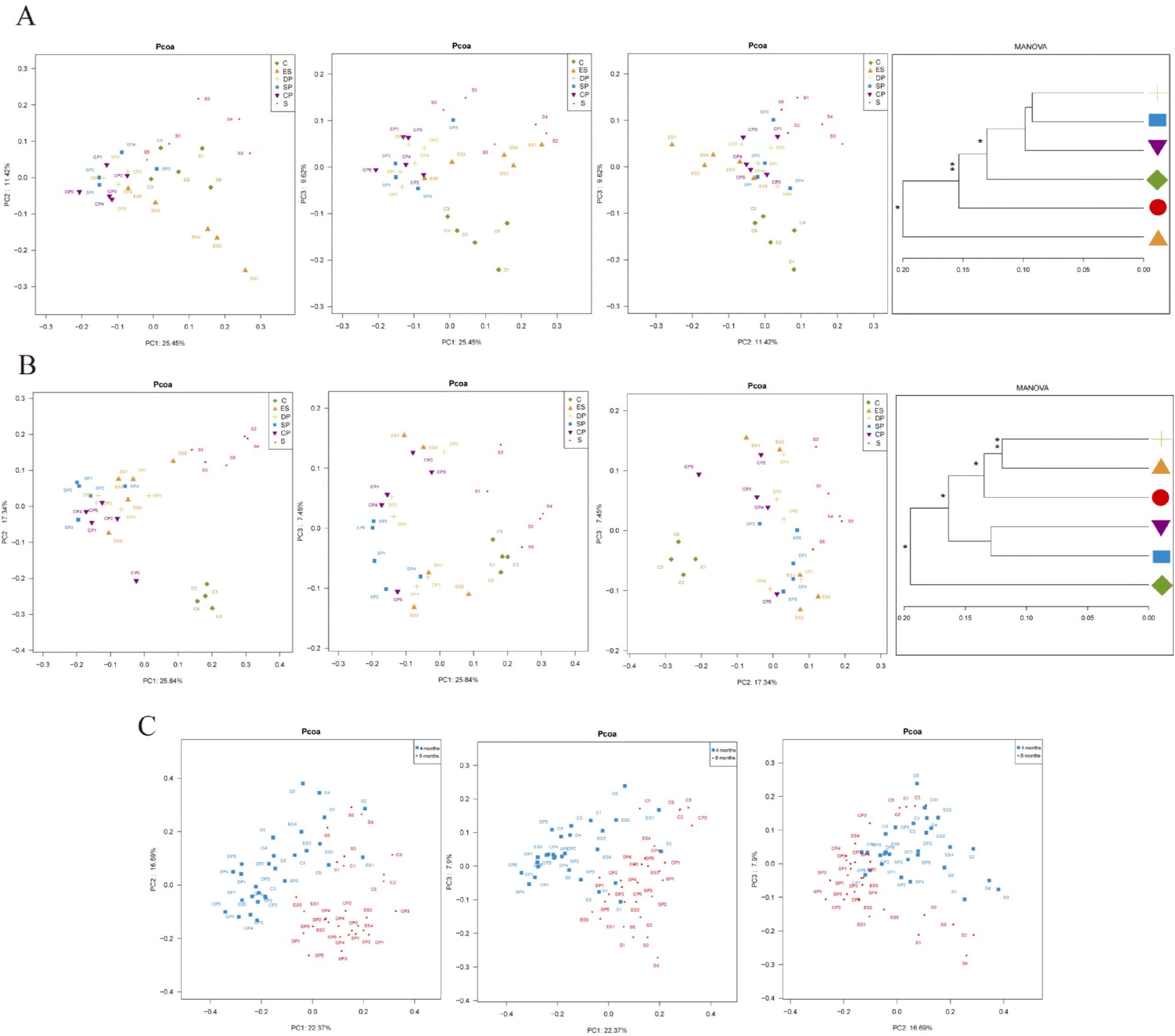
Principal coordinate analysis (PCoA) and clustering analysis. (A) 4 months. (B) 8 months. (C) Two points at 4 and 8 months. Note: the MANOVA significance was also indicated: * *P* <0.05; ** *P* < 0.01. C, casein; ES, emulsion-type sausage; DP, dry-cured pork; SP, stewed pork; CP, cooked pork; S, soy.

#### Composition of gut microbiota

On the phylum level, Bacteroidetes and Firmicutes were the predominant phyla (Fig. 2A and B). Hierarchical clustering analysis indicated that the SPD group was different from other groups at 4 months, but the dry-cured pork protein diet (DPPD) and CPPD groups revealed a significant difference from other groups at 8 months in microbial composition. Furthermore, Bacteroidetes abundance increased but Verrucomicrobia abundance markedly decreased during feeding (Fig. 2C).

**Figure 2.**
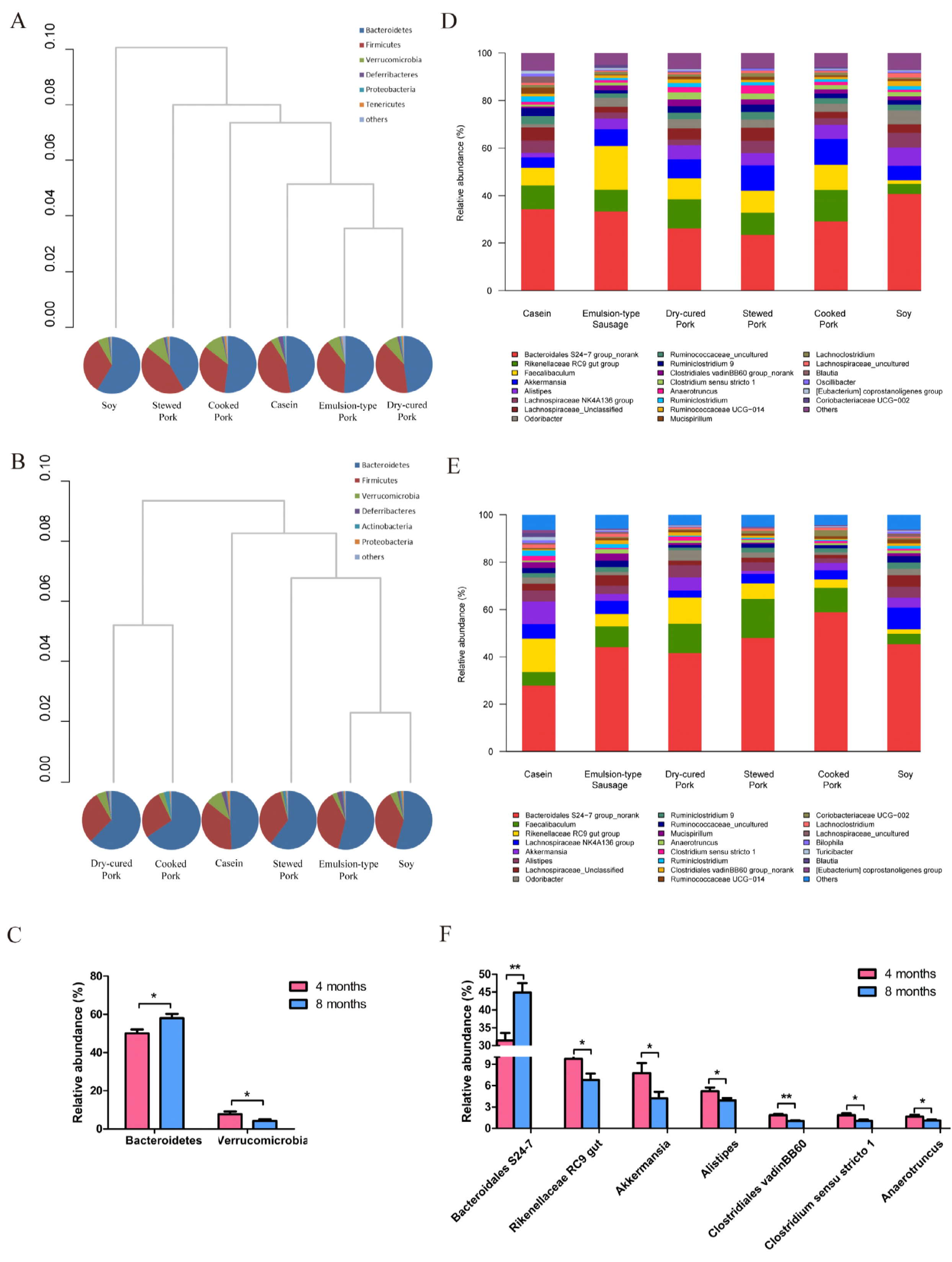
Composition of gut microbiota. (A) The phylum-level taxonomic composition of fecal microbiota at 4 months. (B) The phylum-level taxonomic composition of fecal microbiota at 8 months. (C) Effects of feeding time on the phylum level abundance of 16S rRNA gene sequences. (D) The genus-level taxonomic composition of fecal microbiota at 4 months. (E) The genus-level taxonomic composition of fecal microbiota at 8 months. (F) Effects of feeding time on the genus level abundance of 16S rRNA gene sequences. Note: pie charts show the composition of fecal microbiota at the phylum level. Bray–Curtis similarity cluster analysis shows the similarity and difference of microbial composition in multiple samples. The significance is also indicated: * *P* < 0.05; ** *P* < 0.01.

On the genus level, Bacteroidales S24-7 was the most abundant genus at 4 months, accounting for 31.43% of the fecal microbial population, followed by *Rikenellaceae RC9 gut* (9.75%). At 8 months, Bacteroidales S24-7 and *Faecalibaculum* were the most prevalent genera, accounting for 44.88% and 9.81% of the total count in all diet groups, respectively (Fig. 2D and E). Moreover, seven species showed time-dependent changes. The abundance of Bacteroidales S24-7 increased from 4 to 8 months, whereas those of *Rikenellaceae RC9 gut, Akkermansia, Alistipes,* Clostridiales vadinBB60, *Clostridium sensustricto 1* and *Anaerotruncus* were dramatically reduced (Fig. 2F).

Further analysis revealed that eight of the top twenty dominant genera had significantly changed after 4 and 8 months (Fig. 3A and B). At 4 months, the SPD group had lower abundance of *Rikenellaceae RC9 gut* than the CPPD, DPPD and CD groups, and lower abundance of *Faecalibaculum* than the ESPD and CPPD groups. However, the SPD group had higher abundance of Lachnospiraceae than the DPPD, ESPD and CPPD groups. The CD significantly decreased the abundances of *Alistipes* and Clostridiales vadinBB60 compared with other diets, whereas it increased the abundances of *Blautia* and *Oscillibacter*. In the meat protein diet groups, *Anaerotruncus* was specifically higher in the SPPD group, with Clostridiales vadinBB60 having increased significantly in the DPPD group and *Faecalibaculum* in the ESPD group.

**Figure 3.**
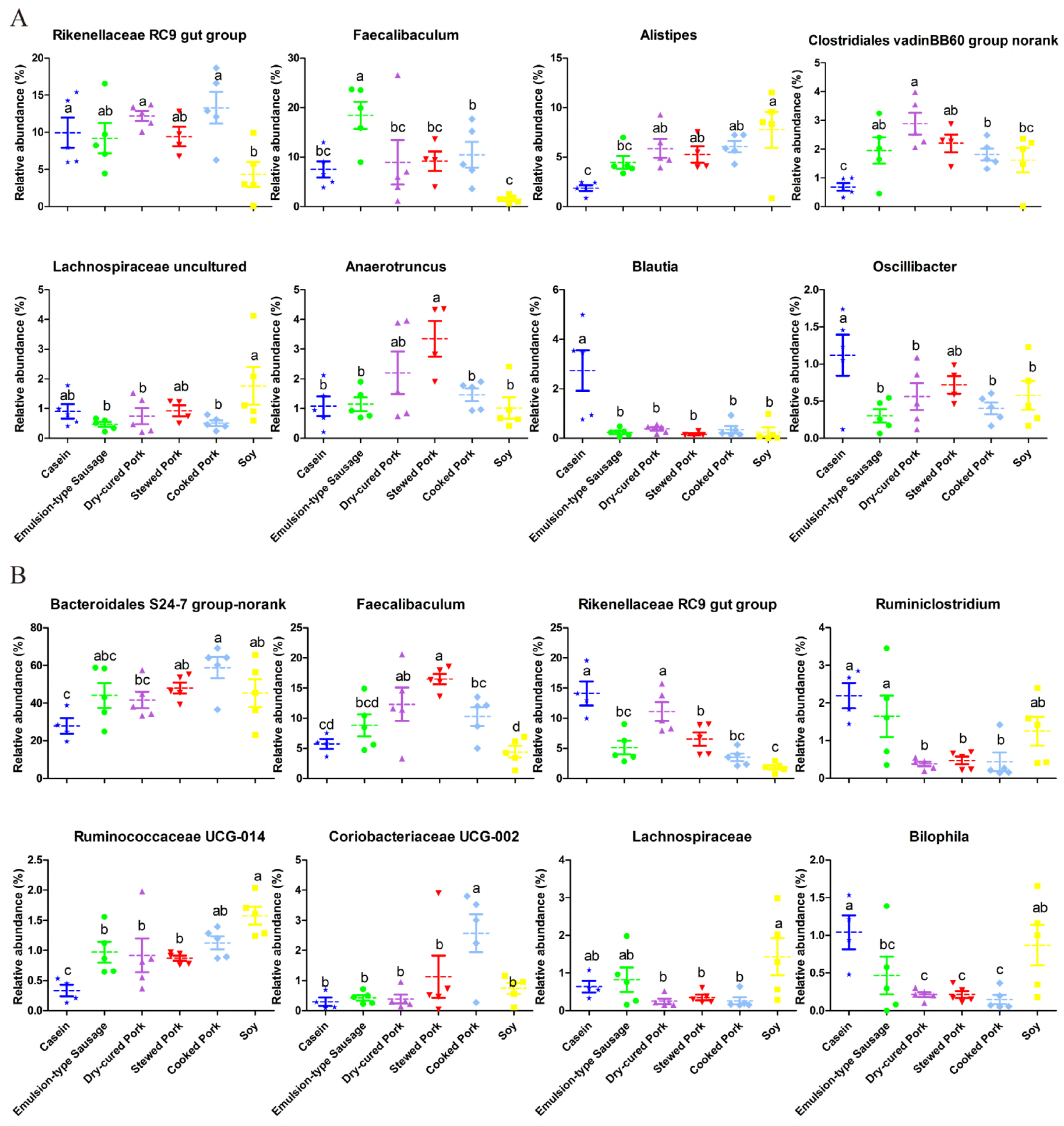
Effects of different protein diets on the top 20 microbial genera. (A) Microbial relative abundance in response to six protein diets at 4 months. (B) Microbial relative abundance in response to six protein diets at 8 months. Note: the data were analyzed by one-way analysis of variance (ANOVA) and means were compared by the procedure of Duncan’s multiple-rangecomparison. The “a, b, c” means with different letters differed significantly (*P* < 0.05), and the microbial genera of significant differences are presented in the figure.

At 8 months, the CD group had lower abundance of Bacteroidales S24-7, *Bilophila* and *Ruminococcaceae UCG-014*, but higher *Rikenellaceae RC9 gut, Ruminiclostridium* and *Faecalibaculum* abundance than the CPPD, DPPD and SPPD groups. The SPD group had lower abundance of *Bilophila* but higher *Faecalibaculum* and Lachnospiraceae abundance than the SPPD, CPPD and DPPD groups. The abundance of *Coriobacteriaceae UCG-002* was the highest for the CPPD group, with *Faecalibaculum* specifically higher in the SPPD group, *Rikenellaceae RC9 gut* in the DPPD group and *Ruminiclostridium* in the ESPD group.

#### Linear discriminant analysis of fecal microbiota

Linear discriminant analysis effect size (LEfSe) analysis revealed 35 different OTUs among the six groups at 4 months (Fig. 4A). OTU36 (*Lachnospiraceae bacterium 609*), OTU23 (Bacteroidales S24-7) and OTU31 (*Eubacterium coprostanoligenes*) were dominant in the CD group. OTU2 (*Faecalibaculum*) and OTU34 (*Coriobacteriaceae UCG-002*) were more abundant in the ESPD group, with OTU35 (Clostridiales vadinBB60) and OTU40 (*Alistipes*) abundance higher in the DPPD group and OTU1 (*Rikenellaceae RC9 gut*) and OTU42 (*Anaeroplasma*) abundance higher in the CPPD group. The SPPD group was abundant in OTU21 (*Lachnospiraceae NK4A136*), OTU39 (*Lachnospiraceae*) and OTU49 (*Clostridium scindens*), which all belong to the Lachnospiraceae family. Eight OTUs that represented the family Bacteroidales S24-7 were more abundant in the SPD group.

**Figure 4.**
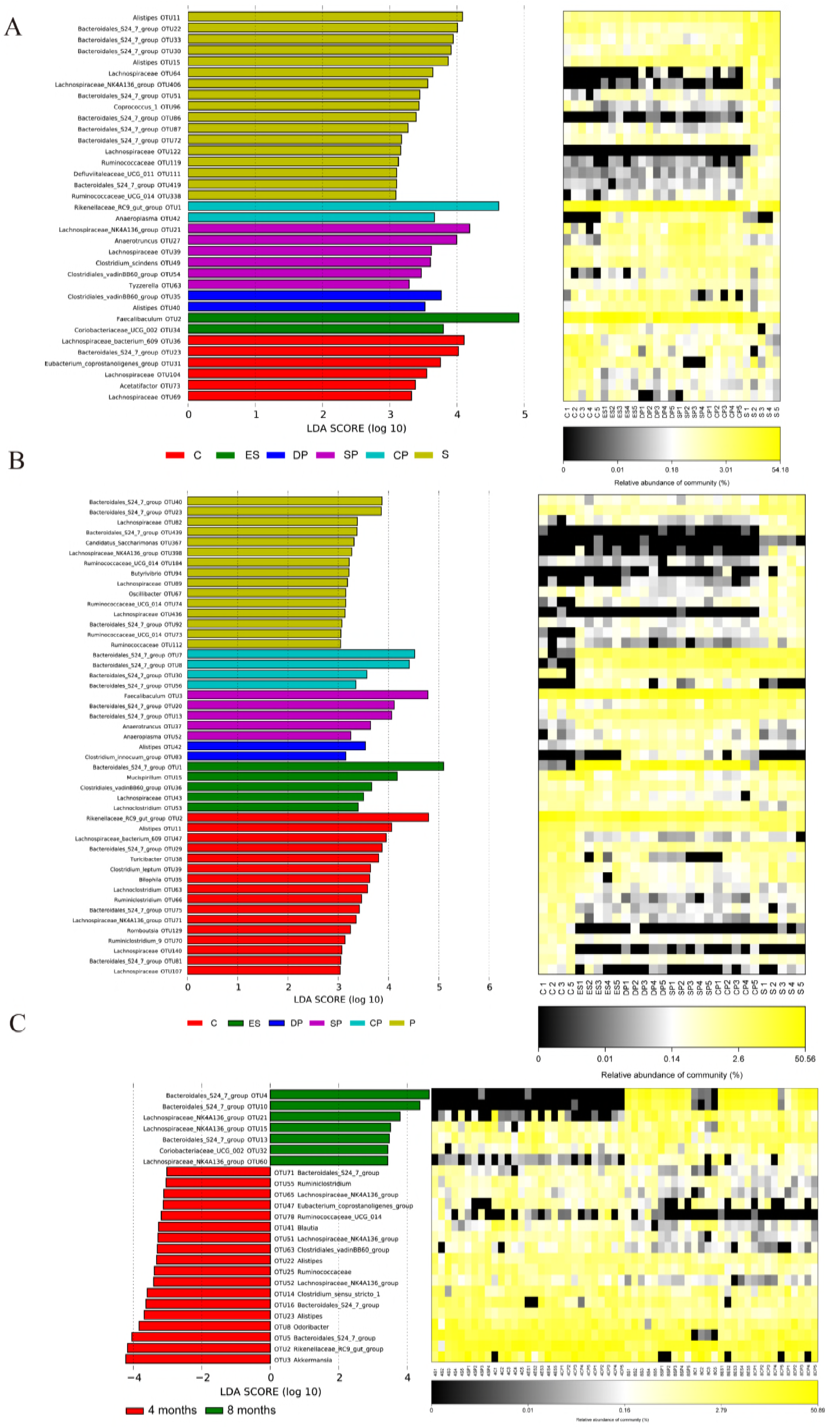
Linear discriminant analysis of fecal microbiota. (A) 4 months. (B) 8 months. (C) Two points at 4 and 8 months. Note: the left histogram shows the LDA scores computed for features at the OTU level. The right heat map shows the relative abundance of OUT (log10-transformed). Each column represents one animal and each row represents the OTU corresponding to the left one. The color intensity scale shows the relative abundance of the OTU (log10-transformed); yellow denotes a high relative abundance of the OTU while black denotes a low relative abundance of the OTU. C, casein; ES, emulsion-type sausage; DP, dry-cured pork; SP, stewed pork; CP, cooked pork; S, soy.

Nevertheless, 47 different OTUs were observed at 8 months (Fig. 4B). The CD group was enriched in two and five OTUs that represented the families of Rikenellaceae and Lachnospiraceae, respectively. OTU42 (*Alistipes*) and OTU83 (*Clostridium innocuum*) were more abundant in the DPPD group. OTU1 (Bacteroidales S24-7) and OTU3 (*Faecalibaculum*) were the most dominant in the ESPD and SPPD groups, respectively. Four OTUs that represented the family Bacteroidales S24-7 were more enriched in the CPPD group. Four, five and six OTUs that respectively represented the families of Bacteroidales S24-7, Lachnospiraceae and Ruminococcaceae were more abundant in the SPD group.

The time effect on the composition of microbiota is shown in Fig. 4C. A total of 25 OTUs were found to change over time. Seven of them obviously increased from 4 months to 8 months; these belong to the families of Bacteroidales S24-7 or Lachnospiraceae or the *Coriobacteriaceae UCG-002* genus. The abundances of five and three OTUs that respectively belong to the families of Ruminococcaceae and Rikenellaceae had significantly reduced. Moreover, three OTUs that represented the genera *Odoribacter*, *Akkermansia* and *Bilophila* were also distinctly decreased over feeding time.

#### Functional prediction of microbial genes

The Phylogenetic Investigation of Communities by Reconstruction of Unobserved States (PICRUSt) revealed five differential functions at 4 months, which are associated with carbohydrate metabolism, the endocrine system, neurodegenerative diseases, cancer and the nervous system (Fig. 5A). The ESPD diet upregulated carbohydrate metabolism and nervous system function more than the CD, SPD and SPPD groups, and the meat protein and casein diets caused a downregulation of genes involved in the endocrine system and neurodegenerative diseases compared with the soy protein diet.

**Figure 5.**
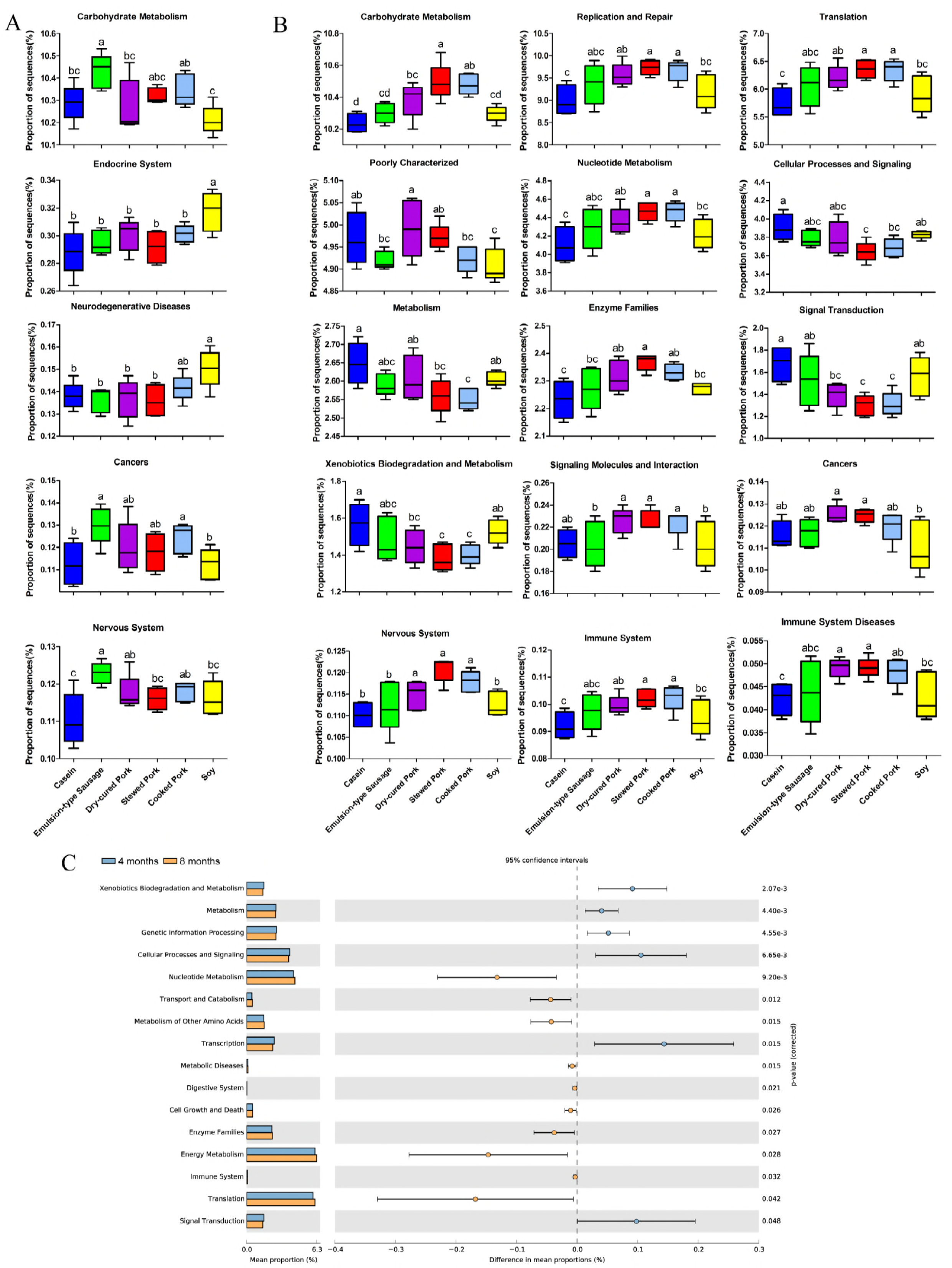
Functional prediction of the microbial genes. (A) 4 months. (B) 8 months. (C) Two points at 4 and 8 months. Note: the data were analyzed by statistical analysis of taxonomic and functional profiles (STAMP) and one-way analysis of variance (ANOVA); means were compared by the procedure of Duncan’s multiple-range comparison. The “a, b, c” means with different letters differed significantly (*P* < 0.05), and the biological function of significant differences are presented in the figure.

At 8 months, 15 gene functions were found to be differentially regulated (Fig. 5B). Compared with the CD group, the SPPD, CPPD and DPPD groups showed a significant reduction in the expression of genes involved in signal transduction, xenobiotics biodegradation and metabolism, while genes involved in carbohydrate metabolism, replication and repair, translation, nucleotide metabolism, enzyme families, the nervous system and the immune system were upregulated. The cellular processes and signaling were significantly downregulated in the SPPD and CPPD groups. In addition, the signaling molecules and interaction were significantly upregulated in the SPPD, CPPD and DPPD groups compared to the SPD group.

In addition, 16 microbial functions were significantly changed during feeding (Fig. 5C). Functions of transcription, cellular processes and signaling, signal transduction, xenobiotics biodegradation, genetic information processing and metabolism were remarkably downregulated. Functions of the immune system, the digestive system, metabolic diseases, cell growth and death, enzyme families, metabolism of other amino acids, transport and catabolism, nucleotide metabolism, energy metabolism and translation were upregulated. This indicates that the diet-induced changes of microbial biological functions are related to the bidirectional communications between the gut microbiota and the host.

### Variations in signaling molecules of the gut–brain axis

To further explore the effects of protein diets on bidirectional communications between the gut and the brain via the peripheral circulatory system, signaling molecules, e.g., peptide YY (PYY), leptin, insulin and serotonin, in serum were quantified. The protein diets significantly affected the concentrations of serotonin, PYY, leptin and insulin in serum at the two time points (Fig. 6A and B).

**Figure 6.**
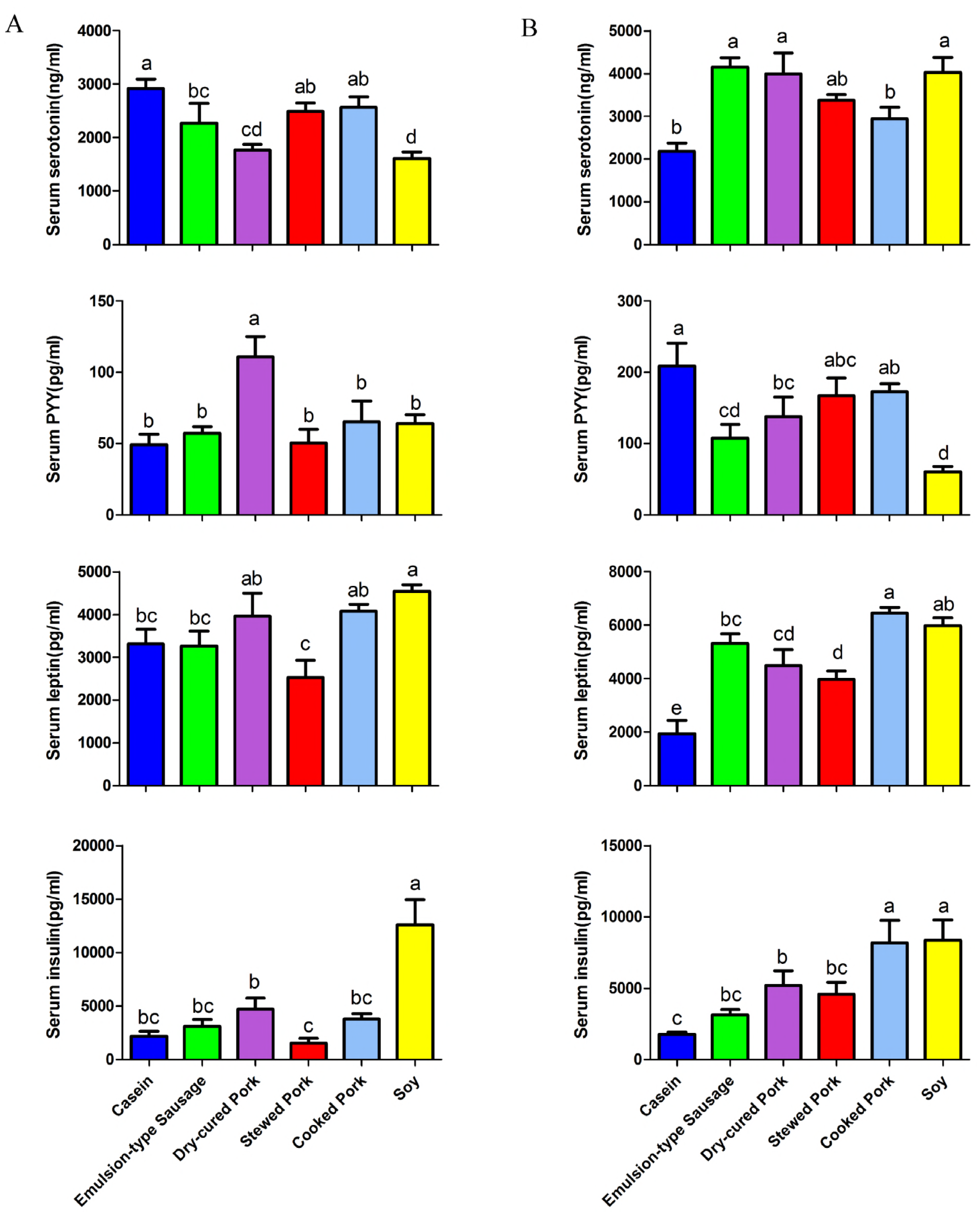
Variations in signaling molecules of the gut–brain axis. (A) 4 months. (B) 8 months. Note: the data were analyzed by one-way analysis of variance (ANOVA) and means were compared by the procedure of Duncan’s multiple-range comparison. The “a, b, c” means with different letters differed significantly (*P* < 0.05).

At 4 months, the concentration of serotonin in the meat protein diet groups was dramatically higher than that in the SPD group, but lower than that in the CD group. Among the meat protein diet groups, the concentration of serotonin was remarkably higher in the CPPD and SPPD groups than in the DPPD group. Nevertheless, the DPPD group had the highest concentration of PYY. On the other hand, the leptin and insulin levels were lower in SPPD group than in the DPPD and SPD groups but did not differ from those in the CD group.

At 8 months, the meat protein diet groups had higher serotonin levels compared with the CD group, and serotonin levels in the CPPD group were lower than in the SPD, ESPD and DPPD groups. As regards PYY, its concentration in the meat protein diet groups was lower than that in the CD group, but higher than that in the SPD group. For meat protein diet groups, the concentration of PYY in the ESPD group was significantly lower than that in the CPPD group. The leptin and insulin levels were lower in the DPPD group than in the SPPD and SPD groups but higher than in the CD group. This indicates that diet-induced changes of gut microbiota could have associations with alterations in signaling molecule levels of the gut–brain axis.

### Key species associated with signaling molecules of the gut–brain axis

To evaluate potential associations between gut microbiota and the signaling molecules of the gut–brain axis, Spearson correlation analysis was performed with dominant OTUs whose relative abundance was at least 0.5% in at least one group. We observed that 28 and 48 OTUs were apparently correlated with signaling molecules including serotonin, PYY, leptin and insulin at 4 and 8 months, respectively (Fig. 7A and B).

**Figure 7.**
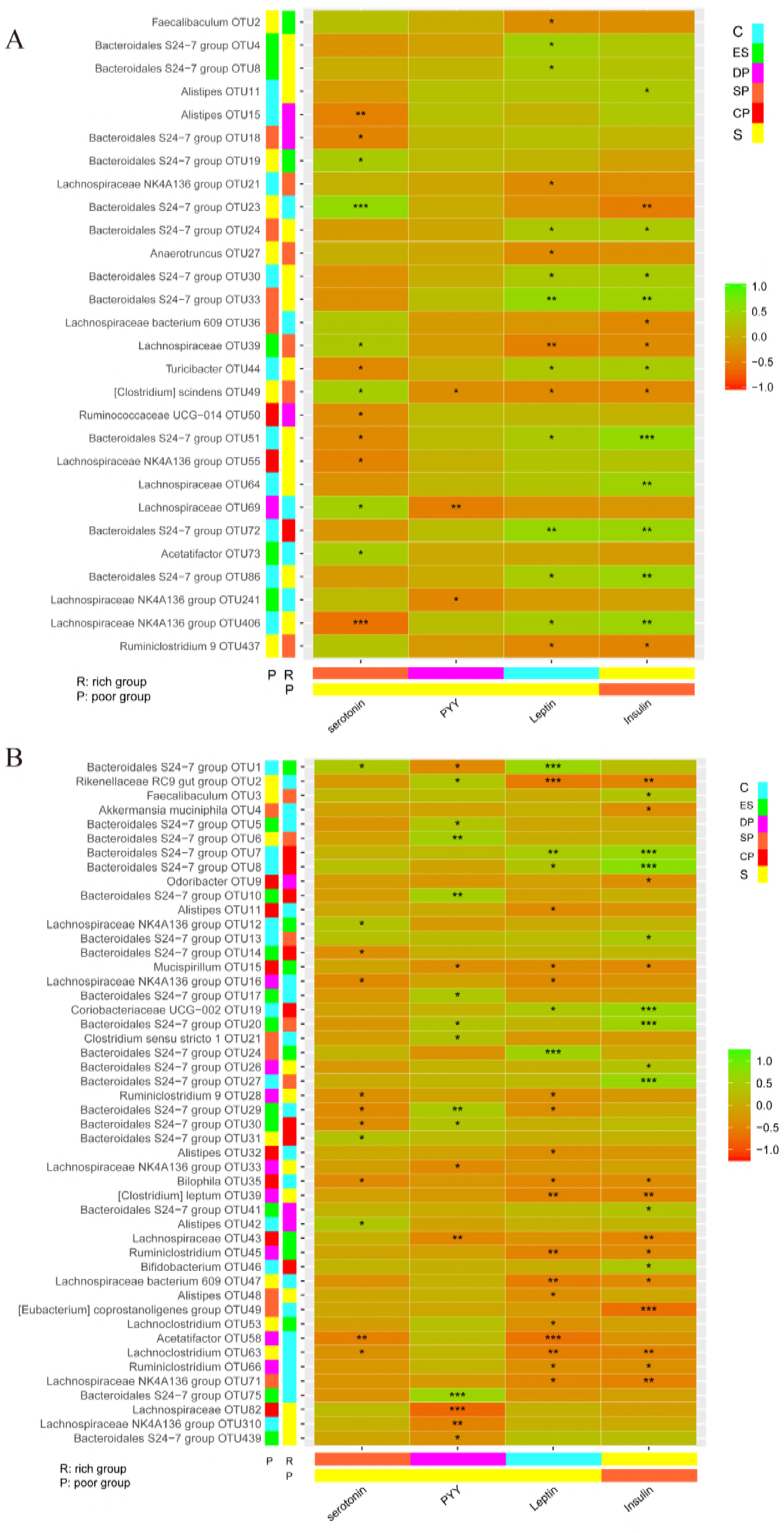
Key species associated with signaling molecules of the gut–brain axis. (A) 4 months. (B) 8 months. Note: each figure has four parts: (1) the large heat map, correlations between microbiota and signaling molecules, where green represents significant positive correlation and red represents significant negative correlation; (2) the bottom bars represent rich/poor groups of signaling molecules; (3) the right bars next to the large heat map represent rich/poor groups of different microbial OTU-level taxa; (4) the independent right color bars depict correlation coefficients between microbiota and signaling molecules. OTUs whose relative abundance is at least 0.5% in at least one group were analyzed; significantly related OTUs are shown in the figures. The significance is also indicated: * *P* <0.05; ** *P* < 0.01; *** *P* < 0.001. C, casein; ES, emulsion-type sausage; DP, dry-cured pork; SP, stewed pork; CP, cooked pork; S, soy; R, rich group; P, poor group.

At 4 months, four and two OTUs that represented the families Lachnospiraceae and Bacteroidales S24-7, respectively, were positively correlated with the concentration of serotonin. However, each of the two OTUs that respectively represented the genus *Lachnospiraceae NK4A136* and the family Bacteroidales S24-7 were all negatively correlated with serotonin levels. Both leptin and insulin levels were positively correlated with six OTUs that represented the family Bacteroidales S24-7, but they were negatively correlated with two OTUs that represented the family Lachnospiraceae. Finally, PYY levels showed a positive statistical relationship with OTU49 (*Clostridium scindens*) and OTU241 (*Lachnospiraceae NK4A136*), which all belong to the family Lachnospiraceae.

At 8 months, OTU1 and OTU31 (Bacteroidales S24-7) and OTU42 (*Alistipes*) were positively correlated with serotonin levels. On the contrary, each of the three OTUs that respectively represented the families Lachnospiraceae and Bacteroidales S24-7 showed a negative correlation with the serotonin levels. Eight OTUs in the families of Bacteroidales S24-7 had a positive correlation with PYY levels, and four OTUs that represented the families of Lachnospiraceae revealed a negative correlation with serotonin levels. The insulin and leptin levels showed a positive correlation with seven and four OTUs, respectively, which all belong to the Bacteroidales S24-7 family. Each of the three OTUs that represented the families of Lachnospiraceae and Ruminococcaceae were also positively correlated with the insulin and leptin levels. Nevertheless, these levels were negatively correlated with OTU2 (*Rikenellaceae RC9 gut*), OTU15 (*Mucispirillum*) and OTU35 (*Bilophila*).

The microbial network analysis of 4-month data indicated nine positive correlations (green lines) and seven negative correlations (red lines) on the OTU level (Fig. 8A). For 8-month data, 47 positive correlations (green lines) and 22 negative correlations (red lines) were observed. (Fig. 8B). *Bilophila* was positively correlated with *Mucispirillum*, Lachnospiraceae and Ruminococcaceae, as was Ruminococcaceae with Lachnospiraceae and *Mucispirillum*; they all showed a negative relationship with Bacteroidales S24-7. However, Bacteroidales S24-7 was positively correlated with Erysipelotrichaceae and Rikenellaceae.

**Figure 8.**
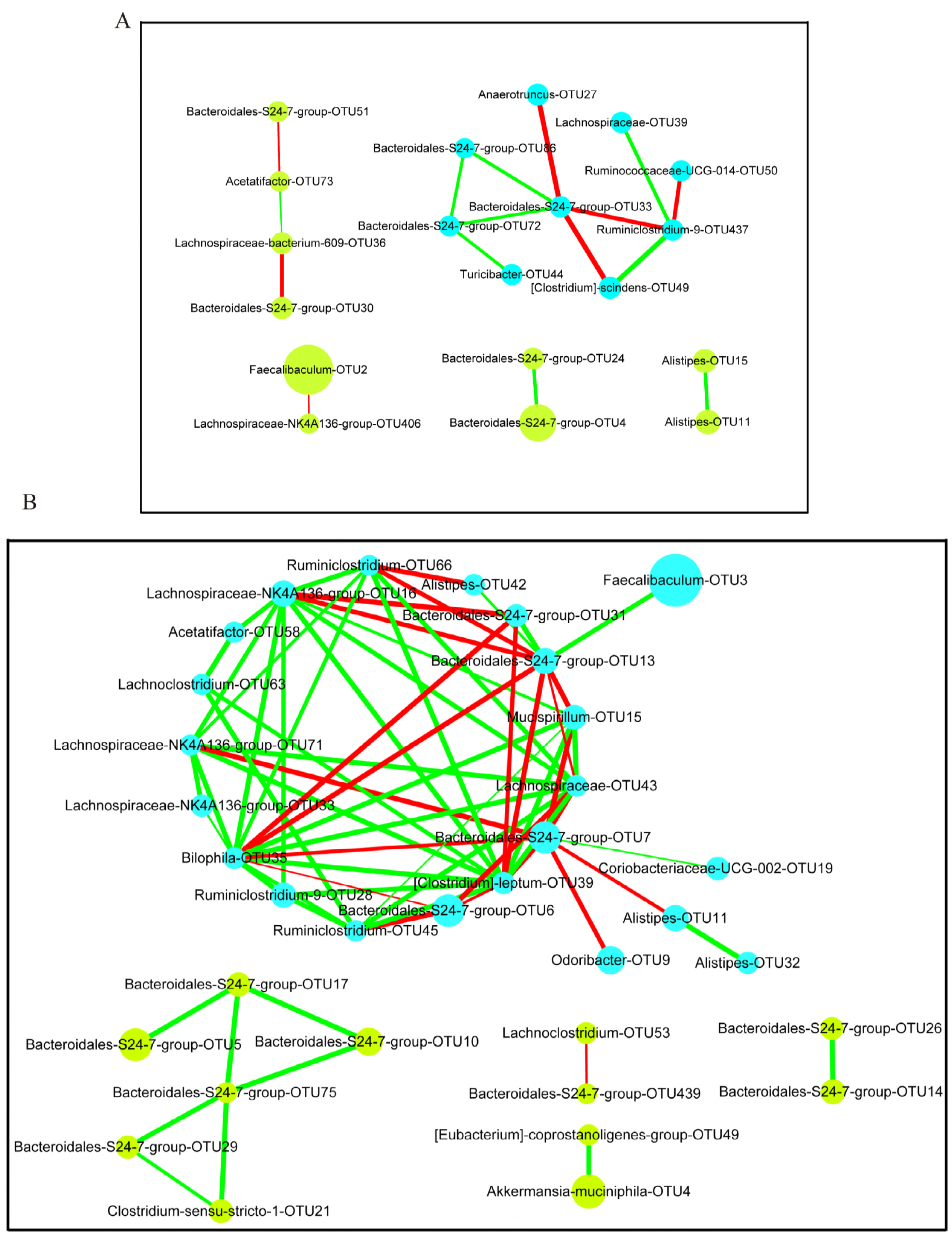
Co-occurrence network of the key species. (A) 4 months. (B) 8 months. Note: nodes represent the OTUs identified by correlation analysis, and the size of the node corresponds to the relative abundance of the OTUs or genera. Each pair of OTUs or genera showing a Spearman correlation coefficient value higher than 0.6 is linked with a connecting line whose thickness corresponds to the coefficient values. The green line represents significant positive correlation while the red line represents significant negative correlation.

## Growth performance

At the baseline (before diet intervention), no significant difference was observed in body weight between any two diet groups. However, the protein diets had a significant impact on the body weight of the mice (Fig. 9A). The body weight of mice in the CPPD, SPPD and ESPD groups increased with feeding time, while the CD induced a great decline in body weight at the 24th week of the experiment. A similar phenomenon was observed in the SPD and DPPD groups at the 30th week of the experiment. At the end of the diet, the body weight of the SPPD group was significantly lower than that of the SPD group, but higher than that of the CD group, which was in line with the average daily gain (ADG). Correspondingly, the average daily feed intake and the feed efficiency were the lowest for the CD group. The average daily feed intake (ADFI) of the SPD group was higher than that of the SPPD group, but no significant difference was observed for feed efficiency (Fig. 9B to D).

**Figure 9.**
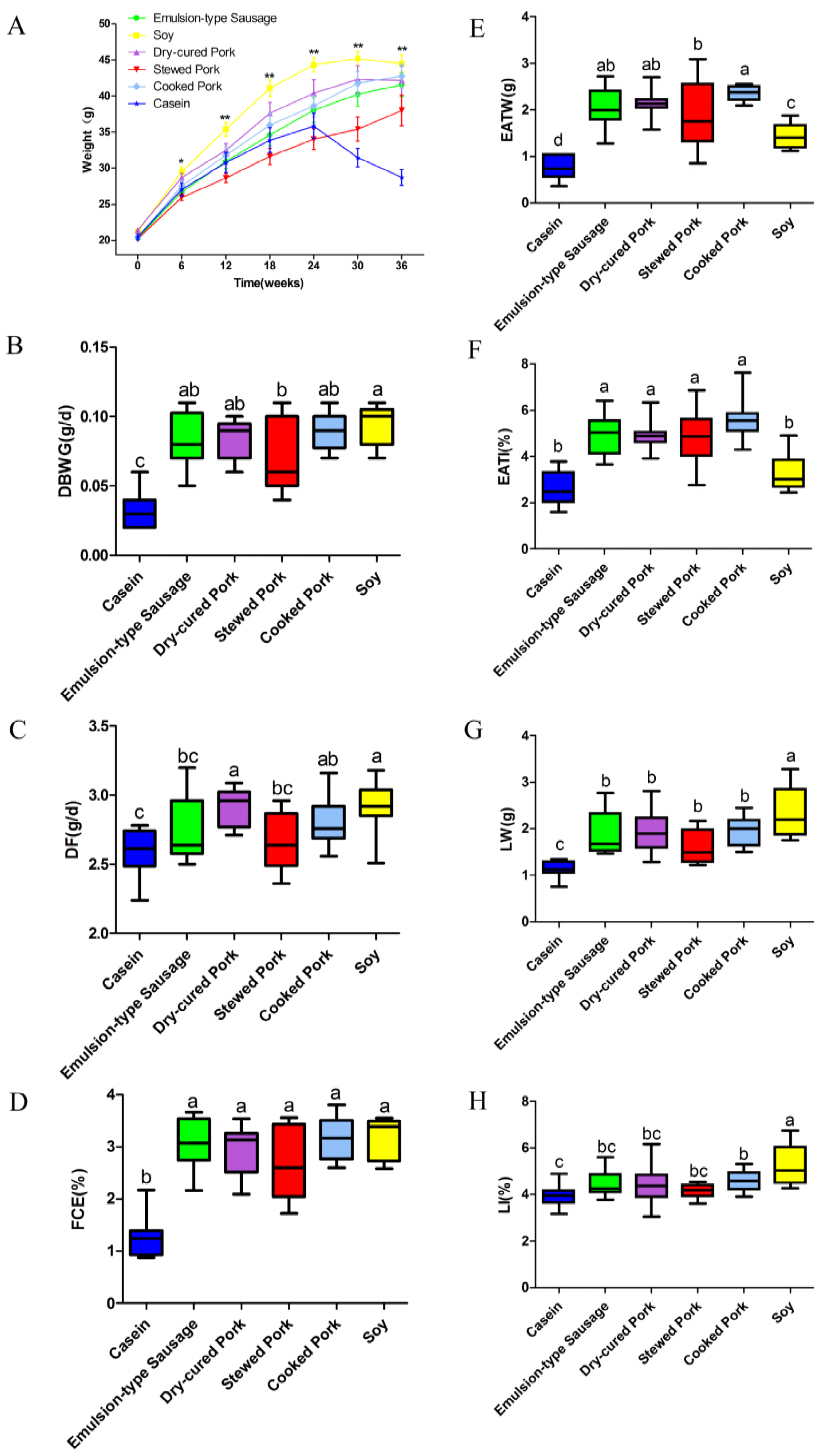
Growth performance. (A) Body weight during feeding. (B) Daily body weight gain. (C) Daily feed intake. (D) Feed conversion efficiency. (E) Absolute weight of epididymaladipose tissue. (F) Relative weight of epididymal adipose tissue to body weight. (G) Absolute weight of liver. (H) Relative weight of liver to body weight. Note: the data at each feeding time point were analyzed by one-way analysis of variance (ANOVA) and means were compared by the procedure of Duncan’s multiple comparison. The asterisks (*) indicate significant differences between diet groups. * *P* < 0.05; ** *P* < 0.01. The “a, b, c” means with different letters differed significantly (*P* < 0.05). ADG, average daily gain; ADFI, average daily feed intake; FCE, feed efficiency; EATW, absolute weight of epididymal adipose tissue; EATI, relative weight of epididymal adipose tissue to body weight; LW, absolute weight of liver; LI, relative weight of liver to body weight.

The development of epididymal adipose tissue and of the liver can reflect, to a certain extent, the body composition of mice as a response to their diet. The protein diets had a distinct impact on the weight of the epididymal adipose tissue and liver (Fig. 9E to H). The non-meat protein diet groups (CD and SPD) had less epididymal adipose than the meat protein diet groups (ESPD, DPPD, SPPD and CPPD). Nevertheless, the liver weights of the meat protein diet groups were lower than those in the SPD group but higher than those in the CD group. This may be related to feed intake and weight gain.

### Discussion

The gut is a complex and dynamic ecosystem. The temporal pattern of microbial survival is the key for finding out core members responding to environmental changes. Many studies have shown that the composition of fecal microbiota is highly correlated with the colonic lumen and mucosa and moderately correlated with the distal small intestine (18). Our previous studies indicated that intake of soy protein, casein and meat proteins altered the composition of cecal microbes in rats (19). In the present study, the diet-induced and temporal changes of microbiota in mice have been analyzed and correlated with signaling molecules of the gut–brain axis.

Obviously, the different protein diets led to different microbial compositions, which may be due to the different digestibility and digested products of processed meat proteins (17) from those of non-meat proteins. It is noteworthy that the responses of gut microbiota to the diet differ between bacterial species; some species responded faster or slower than others to the protein diet.

The microbial structure can be easily affected by the host’s physiological status and by environmental perturbations. However, microbial structure remains stable during development (20). Environmental factors, including diet, may drive them to a new homeostasis. For each genus of which the abundance is affected by the protein diets, all protein diets in this study either increased or decreased its abundance. Similar results have been shown in a short-term, high-fat diet feeding study (10). As is well known, *Akkermansia* is a mucin-dependent bacterium (21) that can stimulate the synthesis and secretion of mucin, which is the main component of the mucus layer and acts as a mucus barrier (22). *Akkermansia* was considered to be a vital biomarker for intestinal health (23) and to aid in the prevention of obesity, diabetes, inflammation and other metabolic disorders (24). Notably, *Akkermansia* was not significantly affected by protein diets at the two time points. However, the abundance of *Akkermansia* was reduced from 4 months to 8 months in the present study. In addition, the Shannon diversity index was also significantly reduced with feeding time. Long-term intake of a certain diet may lead to decreased microbial diversity and to destruction of the ecological balance of gut microbiota. Many studies have shown that low microbial diversity is associated with some metabolic disorders (25, 26). Our microbial network analysis has indicated that 16 and 69 microbial interactions existed in the two time points, respectively. Ruminococcaceae was positively associated with Lachnospiraceae but negatively with Bacteroidales S24-7, which can partly explain the decline in Ruminococcaceae and Lachnospiraceae and the increase in Bacteroidales S24-7.

Dietary modulation can alter the microbial community and metabolic activity (27, 28). Previous studies showed that many members of the Rikenellaceae, Lachnospiraceae and Ruminococcaceae families show a high potential for fermenting dietary proteins (29-31). Short-chain fatty acids (SCFAs) are important microbial metabolites that serve as important nutrients for the gut epithelium and body tissues that can affect the metabolism, immune response and anti-inflammatory function (32, 33). High-protein diets have been shown to affect the production of SCFAs both in human and in rodent models (34, 35). Ruminococcaceae and *Bifidobaterium* are known to be acetate producers. Many members of Lachnospiraceae are capable of producing butyric acid through the fermentation of various substrates (29). In the present study, different protein diets have altered the numbers of SCFA producers. We hypothesize that different dietary proteins affect the levels of SCFAs by changing the composition of the gut microbiota. Some studies showed that SCFAs may affect the production of hormones and neurotransmitters (36, 37). The host–microbe fundamental relationship relies on chemical signaling and nutrient availability (38). Twenty-eight and forty-eight specific OTUs were identified to have distinct correlations with serotonin, PYY, leptin and insulin at 4 and 8 months, respectively. These hormones and neurotransmitters play important roles in the communication between the gut and the brain, especially in terms of appetite and energy balance (39, 40).

Precise regulation of appetite contributes to maintenance of the body’s stable energy metabolism and weight level. Many studies also showed that the gut microbial composition is linked to body weight and average daily gain (41), and that the production of SCFAs can improve the absorptive capacity of the intestine and increase feed efficiency (42). Soy protein isolates were used in obese mice to reduce fat deposition. Similarly, in our study, the non-meat protein diet-fed mice (CD and SPD) had less epididymal adipose tissue than mice fed the meat protein diets (ESPD, DPPD, SPPD and CPPD), even though mice in the SPD group had higher body weights than those in the meat protein diet groups. The liver weights of mice in the meat protein diet groups were lower than in the SPD group but higher than in the CD group; the relative differences are similar to the change in intake and weight gain of the mice. Furthermore, the growth rate of mice fed different protein diets tended to be slow, and even decreased over the feeding period. The growth rates were different from previous studies using a rat model (43), which may be related to the physiological performance of the animals themselves and the time of dietary intervention.

Above all, our results show that specific microbiota dynamically regulate signaling molecule levels, consequently affecting growth performance, suggesting that consuming the same diet for a prolonged time, irrespective of the kind of diet, may adversely affect our health to some extent by altering the microbial composition. Although these results from animal experiments cannot be extrapolated directly to humans, they do provide some evidence and references the composition of a healthy human diet. A healthy diet may help us improve not only gastrointestinal diseases but also other health problems, such as nervous system-related disorders, by regulating the microbial structure and balance. The exact mechanisms remain unclear; more studies are necessary to investigate how diets stably improve health status through re-shaping the gut microbiota composition in the long term.

### Materials and Methods

#### Animals and diets

All experiments were carried out in compliance with the relevant guidelines and regulations of the Ethical Committee of Experimental Animal Center of Nanjing Agricultural University. A total of 60 4-week-old male C57BL/6J mice were obtained from Nanjing Biomedical Research Institute and housed in a specific pathogen-free animal center (SYXK<Jiangsu> 2011-0037). The temperature (20.0 ± 0.5°C) and relative humidity (60 ± 10%) were kept constant during the experiment, with a 12-h light cycle. Mice were fed a standard chow diet during a 2-week acclimation period. Then, animals were assigned to one of six diet groups (ten mice in each group and two per cage), i.e., CD, ESPD, DPPD, SPPD, CPPD and SPD groups. Mice were allowed to access water and diets *ad libitum* for 8 months. Body weights and feed intake of mice were routinely recorded for calculating the ADG and the ADFI. The feed efficiency was expressed as a ratio of ADG to ADFI.

#### Sample collection

After 4 and 8 months of feeding, the feces and blood of mice were collected. The fecal samples from the two mice in the same cages were mixed and stored at −80°C for the microbial composition analysis. Blood samples were centrifuged at 12,000 g for 30 min to pellet the blood cells and serum samples were stored at −80°C. After 8 months, all the mice were euthanized by cervical dislocation, and the epididymal adipose and liver tissues were taken and weighed. Relative weights of epididymal adipose and liver tissues were calculated according to body weight.

#### Serum signaling molecules of the gut–brain axis

The serum signaling molecules of the gut–brain axis, including peptide YY (PYY), leptin and insulin, were measured using the Milliplex magnetic bead mouse metabolic hormone multiplex panel (MMHMAG-44K; EMD-Millipore, Billerica MA), and serotonin (5-hydroxytryptamine, 5-HT) was quantified using a serotonin ELISA kit (KA2518, Abnova, USA) according to the manufacturer’s protocols.

#### 16S rRNA gene sequencing

Total genomic DNA in fecal samples was extracted using the QIAamp DNA Stool Mini Kit (No. 51504, Qiagen, Germany) according to the manufacturer’s instructions. The DNA was quantified by a Nanodrop^®^ spectrophotometer (Nanodrop2000, Thermo, USA, Shanghai). Purified DNA was used to amplify the V4 region of 16S rRNA, which is associated with the lowest taxonomic assignment error rate (44). Polymerase chain reaction (PCR) was performed in triplicate. Amplicons were extracted, purified and quantified. The pooled DNA product was used to construct Illumina Pair-End library following Illumina’s genomic DNA library preparation procedure. Then the amplicon library was paired-end sequenced (2 × 250) on an IlluminaMiSeq platform (Shanghai Biozeron Co., Ltd) according to the standard protocols.

#### Bioinformatics analysis

Raw fastq files were trimmed and chimeric sequences were identified and removed from all samples to reduce noise, and operational taxonomic units (OTUs) were clustered with ≥97% similarity. In line with the results of the OTU clustering analysis, we can define the relative abundance of each OTU at different taxonomic levels and carry out a variety of diversity index analyses. Community richness estimator (Chao and ACE), diversity indices (Shannon and Simpson), and Good’s coverage were calculated (45). Principal coordinate analysis (PCoA) and clustering analysis were applied on the basis of the OTUs to offer an overview of the fecal microbial composition (46). Multivariate analysis of variance (MANOVA) analysis was conducted to further confirm the observed differences. LEfSe analysis was carried out to discover biomarkers for fecal bacteria and to distinguish between biological conditions among different groups (47). Besides, the Spearman correlation coefficients were assessed to determine the relationships between microbiota and signaling molecules of the gut–brain axis.

#### Functional prediction of the microbial genes

PICRUSt program based on the Kyoto Encyclopedia of Genes and Genomes (KEGG) database was used to predict the functional alteration of fecal microbiota in different samples (48). The OTU data obtained were used to generate BIOM files formatted as input for PICRUSt v1.1.09 with the make.biom script usable in the Mothur. OTU abundances were mapped to Greengenes OTU IDs as input to speculate about the functional alteration of microbiota.

#### Statistical analysis

The diet effect was evaluated by one-way ANOVA with SAS software (SAS Institute Inc., Cary, NC, USA). Means were compared and the significance threshold was set at 0.05 for statistical analyses. Figures were constructed using the GraphPad Prism (version 5.0.3, San Diego, CA, USA).

The details are described in supplementary file, DOCX file, 40.7 KB.

## Acknowledgements

This study was financially supported by grants from National Natural Science Foundation of China (No. 31530054). We would like to thank Jiangsu Department of Education (PAPD) for support and LetPub (www.letpub.com) for providing linguistic assistance during the preparation of this manuscript.

